# Neural responses to naturalistic clips of behaving animals in two different task contexts

**DOI:** 10.1101/268144

**Authors:** Samuel A. Nastase, Yaroslav O. Halchenko, Andrew C. Connolly, M. Ida Gobbini, James V. Haxby

## Abstract

Neuroimaging studies of object and action representation often use controlled stimuli and implicitly assume that the relevant neural representational spaces are fixed and context-invariant. Here we present functional MRI data measured while participants freely viewed brief naturalistic video clips of behaving animals in two different task contexts. Participants performed a 1-back category repetition detection task requiring them to attend to either animal taxonomy or animal behavior. The data and analysis code are freely available, and have been curated according to the Brain Imaging Data Structure (BIDS) standard. We thoroughly describe the data, provide quality control metrics, and perform a searchlight classification analysis to demonstrate the potential utility of the data. These data are intended to provide a test bed for investigating how task demands alter the neural representation of complex stimuli and their semantic qualities.

---

The human brain rapidly deploys semantic information during perception to facilitate our interaction with the world. These semantic representations are encoded in the activity of distributed populations of neurons (Haxby et al., 2001; Kriegeskorte et al., 2008b; McClelland and Rogers, 2003) and command widespread cortical real estate (Binder et al., 2009; Huth et al., 2012). The neural representation of a stimulus can be described as a location (i.e., response vector) in a high-dimensional neural representational space (Haxby et al., 2014; Kriegeskorte and Kievit, 2013). This resonates with behavioral and theoretical work describing mental representations of objects and actions as being organized in a multidimensional psychological space (Attneave, 1950; Edelman, 1998; Gärdenfors and Warglien, 2012; Shepard, 1958, 1987). Current applications of this framework to neural representation (e.g., Kriegeskorte et al., 2008b) often implicitly assume that these neural representational spaces are relatively fixed and context-invariant. In contrast, earlier work emphasized the importance of attention and task demands in actively reshaping representational space (Kruschke, 1992; Nosofsky, 1986; Shepard, 1964; Tversky, 1977).

Here we present functional MRI data measured while participants freely viewed brief naturalistic video clips of animals behaving in their natural environments (Nastase et al., 2017). Participants performed a 1-back category repetition detection task requiring them to attend to either animal behavior or taxonomy. There are several benefits to using dynamic, naturalistic stimuli. They convey rich perceptual and semantic information (Bartels and Zeki, 2004; Huth et al., 2012) and more fully sample neural representational space than conventional stimuli (Haxby et al., 2014). Furthermore, natural vision paradigms have greater ecological validity (Felsen and Dan, 2005), and dynamic stimuli have been shown to drive reliable neural responses across individuals (Hasson et al., 2010; Haxby et al., 2011). These data are intended to provide a test bed for investigating how task demands alter the neural representation of complex stimuli and their semantic qualities.

Twelve right-handed adults (seven female; mean age = 25.4 years, SD = 2.6, range = 21-31) with normal or corrected-to-normal vision were sampled from the Dartmouth College community to participate in the experiment. Participants reported no history of psychiatric or neurological disorders. All participants provided written, informed consent prior to participating in the study in compliance with the Committee for the Protection of Human Subjects at Dartmouth College, including a provision for data to be shared with other researchers around the world or on a publicly available data archive. The study was approved by the Institutional Review Board of Dartmouth College, and participants received monetary compensation for their participation. All data were collected between June 1 and September 6, 2013.

We implemented a full factorial repeated measures design (Fisher, 1935) comprising five taxonomic categories, four behavioral categories, and two tasks. The five taxonomic categories were primates, ungulates, birds, reptiles, and insects. The four behavioral categories were eating, fighting, running, and swimming. Crossing the taxonomy and behavior factors yielded 20 total taxonomy–behavior conditions. The animal taxonomy (i.e., object/form category) and behavior (i.e., action/motion category) factors were chosen as these are thought to rely on somewhat distinct, relatively well-studied neural pathways (Connolly et al., 2012; Giese and Poggio, 2003; Oosterhof et al., 2013; Sha et al., 2015; Wurm et al., 2017). The taxonomic and behavioral categories roughly correspond to intermediate levels of noun and verb hierarchies (Fellbaum, 1990; Rosch, 1975). We designed the experiment under the assumption that the stimulus dimensions conveying taxonomic and behavioral information are not integral (i.e., producing facilitation or interference across factors; Garner and Felfoldy, 1970). However, this may not hold in practice; for example, some taxonomic features may be necessary for behavior categorization, and certain taxa (e.g., birds) may interfere with the recognition of certain behaviors (e.g., running). While the taxonomy and behavior factors are fully crossed at the category level, it is not feasible to orthogonalize lower-level correlates (e.g., motion energy, the specific animal performing each action) in natural vision paradigms.

Each of the 20 taxonomy–behavior conditions comprised two unique 2 s video clips, as well as horizontally flipped versions of each clip for 80 visually unique stimuli in total. Video clip stimuli were sampled from nature documentaries (Life, Life of Mammals, Microcosmos, Planet Earth) and high-resolution YouTube videos. Video clips were edited using the free FFmpeg software package for handling multimedia files (https://www.ffmpeg.org). Stimuli were back-projected onto a screen located at the back of the scanner bore using a Panasonic PT-D4000U projector and viewed via a mirror mounted on the head coil. Video clips subtended a visual angle of ~16.5 degrees horizontally and ~11 degrees vertically. Stimuli were presented using PsychoPy (v1.76.00; http://www.psychopy.org; Peirce, 2007).

In designing the experiment we adopted a condition-rich ungrouped-events design (Kriegeskorte et al., 2008a). Each trial consisted of a 2 s video clip presented without sound followed by a 2 s fixation period for a trial onset asynchrony of 4 s. Each of the 80 stimuli was presented once each run. This type of design has been argued to be particularly efficient for characterizing the pairwise distances between neural response patterns (Aguirre, 2007; Kriegeskorte et al., 2008a). When convolved with a hemodynamic response function, this design matrix will yield highly overlapping response predictors. The response magnitude for each condition can be recovered using a conventional regression model (e.g., Nastase et al., 2017), or regularized regression can be used to predict responses based on an explicit model of stimulus features (e.g., Nishimoto et al., 2011). Each of the 80 unique stimuli can be treated as a separate condition (Kriegeskorte et al., 2008a), or 20 conditions can be defined at the category level by collapsing across the four exemplar clips per taxonomy–behavior condition (Nastase et al., 2017).

In addition to the 80 stimuli, each run included four taxonomy repetition events, four behavior repetition events, and four null fixation events. This resulted in 92 events per run, plus an additional 12 s fixation appended to the beginning and end of each run, for a total run duration of 392 s (~6.5 min). Ten unique runs were created and run order was counterbalanced across participants using a Latin square (Fisher, 1935). Each run was constructed in the following way. First, a pseudorandom trial order containing all 80 stimuli and no taxonomic or behavioral category repetitions was assembled. Second, eight additional stimuli were inserted at particular locations in the trial order to induce four taxonomic category repetition events and four behavioral category repetitions events. Note that in one run an error occurred where a behavior repetition event was inserted that interrupted a previously inserted taxonomic repetition event; this error went unnoticed during data collection but is explicitly noted in text files accompanying the data. These sparse repetition events were inserted such that a repetition event of both types occurred within each quarter of the run. We ensured that the same clip exemplar (or the horizontally mirrored version) never occurred twice consecutively, and that for each taxonomic or behavioral category repetition, the repetition stimulus varied along the other dimension. Finally, four 2 s null events comprising only a fixation cross were inserted at pseudorandom locations in the trial order to effect temporal jittering. One of the four null fixation events occurred each quarter of the run and did not interrupt repetition events. This resulted in an overall scan duration of ~65 min.

Prior to scanning, participants were verbally familiarized with the task and the categories. At the beginning of each run, participants received written instructions indicating that they should pay attention to either taxonomy or behavior and press the button only when they observed a category repetition of that type. Participants were informed that they should ignore repetitions of the unattended type during that run. Button presses were only required for the sparse repetition events (not for non-repetitions) and the same button was used for repetitions of both types. Although responses were collected for repetition events to ensure task compliance, this task was not intended to robustly measure response latencies. We use the term attention loosely here, as performing the 1-back category repetition detection task also requires categorization, working memory, and motor processes. Participants were instructed to maintain fixation only during the fixation periods, and freely viewed the video clip stimuli (cf. Çukur et al., 2013). Behavioral responses for repetition events were collected using a single two-button Lumina LS-PAIR response pad (Cedrus, San Pedro, CA) held in the right hand.

All functional and structural images were acquired using a 3 T Philips Intera Achieva MRI scanner (Philips Healthcare, Bothell, WA) with a 32-channel phased-array head coil. Functional, blood-oxygenation-level-dependent (BOLD) images were acquired in an interleaved fashion using gradient-echo echo-planar imaging with a SENSE parallel imaging factor of 2 (Pruessmann et al., 1999): TR/TE = 2000/35 ms, flip angle = 90°, resolution = 3 mm^3^ isotropic, matrix size = 80 × 80, FoV = 240 × 240 mm, 42 transverse slices with full brain coverage and no gap. At the beginning of each run, two dummy scans were acquired to allow for signal stabilization. Ten runs were collected for each participant, each consisting of 196 functional volumes totaling 392 s (~6.5 min) in duration. At the end of each session, a T1-weighted structural scan was acquired using a high-resolution single-shot MPRAGE sequence: TR/TE = 8.2/3.7 ms, flip angle = 8°, resolution = 0.9375 × 0.9375 × 1.0 mm^3^ voxels, matrix size = 256 × 256, FoV = 240 × 240 × 220 mm^3^. The BOLD signal reflects metabolic demands and serves as a rough proxy for neural activity (primarily local field potentials; Logothetis et al., 2001).

All data have been curated and organized according to the Brain Imaging Data Structure (BIDS) standards (Gorgolewski et al., 2016), and are freely available via the OpenNeuro repository (https://openneuro.org; Poldrack and Gorgolewski, 2017). Data are version-controlled and conveniently accessible using the DataLad data distribution (http://datalad.org; Halchenko et al., 2017) from their original location at http://datasets.datalad.org/?dir=/labs/haxby/attention, as well as from OpenNeuro at https://openneuro.org/datasets/ds001087/versions/00002 and OpenfMRI at https://openfmri.org/dataset/ds000233. According to the BIDS conventions, data are stored in separate directories for each participant alongside the scripts used to compile the data, a descriptive text file, and a tab-separated text file describing participant demographics. Within each participant’s directory, anatomical and functional images are stored in separate directories. Both anatomical and functional images are stored in compressed Neuroinformatics Informatics Technology Initiative (NIfTI-1) format (Cox et al., 2003). Structural images were de-faced for anonymization purposes using an automated masking procedure (Hanke et al., 2014). Each functional run is accompanied by a file describing the acquisition parameters as well as a tab-separated text file describing following for each event: the filename of the clip stimulus, the onset time, duration (2 s), taxonomy–behavior condition, taxonomic category, and behavioral category of the stimulus, as well as whether the stimulus was horizontally mirrored, and whether the event was a repetition or not (and of what type). Participant-specific button presses and their associated response times are also included in the table. Derived data, resulting from preprocessing or other analyses are stored separately in the top-level directory and recapitulate a similar directory structure.

Behaviorally, participants reported category repetitions with high accuracy (99% for both tasks, as reported in Nastase et al., 2017). Although this suggests that participants allocated attention sufficiently to perform the task, it precludes investigators from relating the magnitude of attentional demands to neural responses. We did not design the experiment with a “baseline” or “no task” condition, as it is unclear what this would entail in the context of natural vision paradigms, and any claims about task demands must rely on relative differences between the two tasks.

Organizing data in the standardized BIDS format facilitates the use of portable analysis tools called BIDS Apps (Gorgolewski et al., 2017). To assess the general quality of the data, we used the MRIQC tool (v0.9.6; https://github.com/poldracklab/mriqc; Esteban et al., 2017). Across all participants and runs, median temporal signal-to-noise ratio (tSNR) was 64.73 (range: 31.06-89.14), which approximates the expected tSNR given 3 mm isotropic voxels and 3 T magnetic field strength (Triantafyllou et al., 2005), and is comparable to existing data sets (e.g., Sengupta et al., 2016). Mean framewise displacement (Power et al., 2012) was on average 0.15 mm (range: 0.10-0.44 mm) across participants and runs, indicating fairly low head motion.

To verify that events were annotated correctly, we performed a simple multivariate analysis. Data were first preprocessed using the fmriprep BIDS App (v1.0.0-rc5; https://github.com/poldracklab/fmriprep; Esteban et al., 2017), a Nipype-based tool (Gorgolewski et al., 2011). Cortical surfaces were reconstructed from anatomical scans using FreeSurfer (v6.0.0; https://surfer.nmr.mgh.harvard.edu; Dale et al., 1999) and spatially normalized to the fsaverage6 template based on sulcal curvature (Fischl et al., 1999). Functional images were corrected for slice-timing (Cox, 1996), head motion (Jenkinson et al., 2002), and aligned to the anatomical image (Greve and Fischl, 2009). Functional data were not explicitly spatially smoothed. We then used a general linear model implemented in AFNI (v17.1.02; https://afni.nimh.nih.gov; Cox, 1996) to estimate response patterns for the 20 taxonomy-behavior conditions in each run per task. Nuisance regressors comprised framewise displacement (Power et al., 2012), the first six principal components from an automatic anatomical segmentation of cerebrospinal fluid (aCompCor; Behzadi et al., 2007; Zhang et al., 2001), and de-meaned head motion parameters and their derivatives, regressors for repetition events and button presses, as well as first-through third-order Legendre polynomials.

We then used linear support vector machines (SVMs; Boser et al., 1992; Chang and Lin, 2011) in surface-based searchlights (10 mm radius; Kriegeskorte et al., 2006; Oosterhof et al., 2011) to classify taxonomic and behavioral categories. We used a leave-one-category-out cross-classification approach in both cases: to classify the five taxonomic categories, we trained SVMs on three of the four behavior categories and tested on the left-out behavior category (Figure 2A); to classify the four behavioral categories, we trained SVMs on four of the five taxonomic categories and tested on the left-out taxonomic category (Figure 2B). This approach requires that information about, e.g., behavioral categories, encoded in local response patterns generalizes across both stimuli and taxonomic categories (Kaplan et al., 2015; Nastase et al., 2016; Westfall et al., 2016). All multivariate analyses were performed using PyMVPA (v2.6.3.dev1; http://www.pymvpa.org; Hanke et al., 2009) in the NeuroDebian computational environment (Debian “jessie” 8.5 GNU/Linux with NeuroDebian repositories; http://neuro.debian.net; Hanke and Halchenko, 2011), making heavy use of SciPy (https://www.scipy.org; Jones et al., 2001), NumPy (http://www.numpy.org; Walt et al., 2011), and the IPython interactive shell (https://ipython.org; Perez and Granger, 2007). All scripts used to perform these analyses are provided alongside the data. The resulting searchlight maps corroborate prior work on action and taxonomic category representation (e.g., Connolly et al., 2012; Nastase et al., 2017; Wurm et al., 2017), and demonstrate the potential utility of the data set.

**Figure 1.**
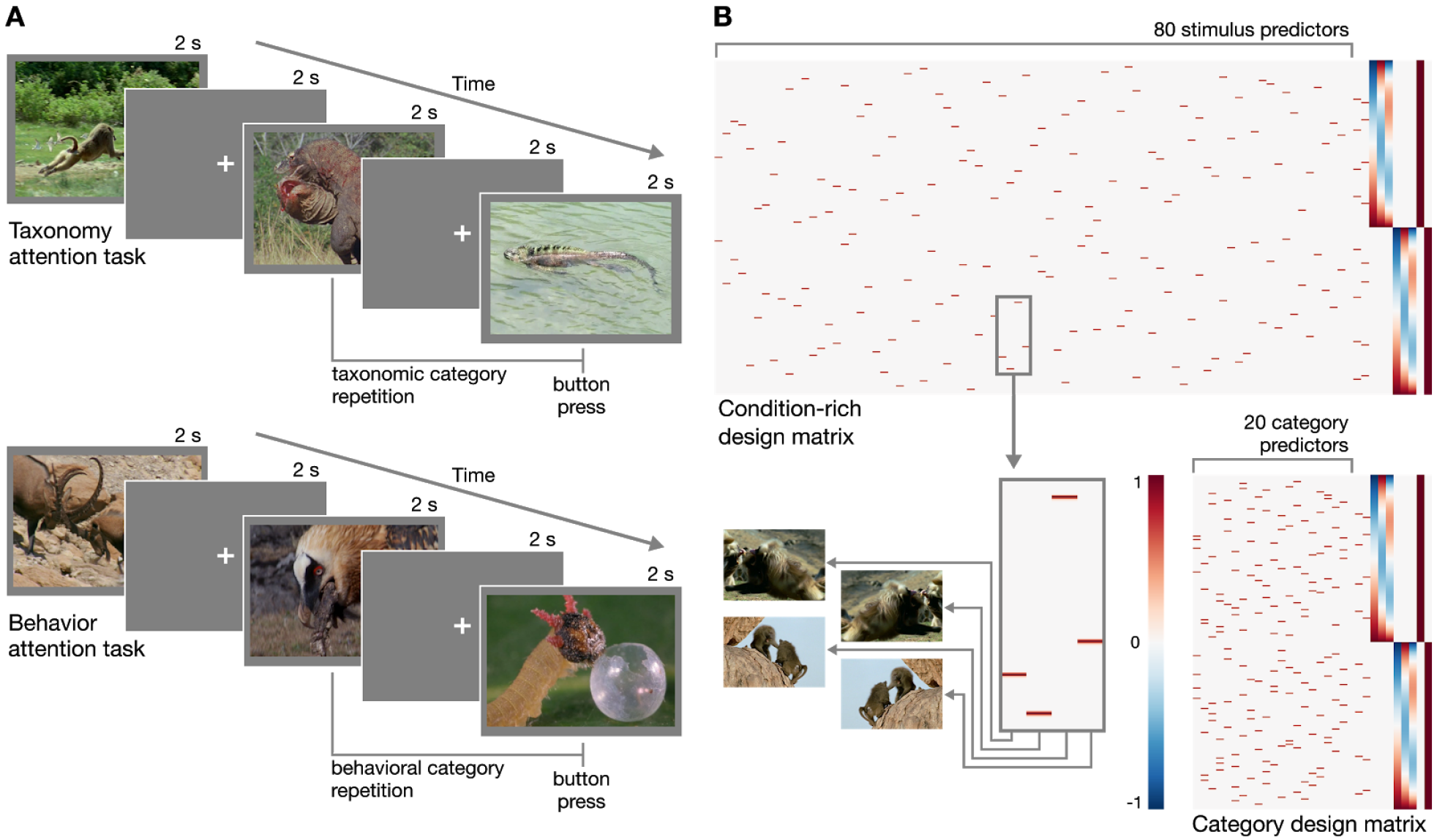
Experimental design. **(A)** Schematic of the rapid event-related design for both taxonomy attention and behavior attention task conditions. In the taxonomy attention task, participants were instructed to press a button if they observed a taxonomic category repetition (e.g., two consecutive clips depicting reptiles; upper). In the behavior attention task, participants were instructed to press a button if they observed a behavioral category repetition (e.g., two consecutive clips depicting animals eating; lower). **(B)** Two example design matrices for predicting hemodynamic responses to the clips over the course of two runs with the taxonomy attention task. In the condition-rich design, each of 80 visually unique stimuli receives a separate predictor (following Kriegeskorte et al., 2008a; upper), while in the category design, the four exemplar clips per taxonomy-behavior condition are collapsed to form 20 category predictors (following Nastase et al., 2017; lower). Hypothesized neural responses are convolved with a simple hemodynamic response function (Cohen, 1997). In this simple example, nuisance regressors for taxonomy and behavior repetition events, first-through third-order Legendre polynomials, and run constants are appended to each design matrix. Figures were created using Matplotlib (https://matplotlib.org; Hunter, 2007) and seaborn (https://seaborn.pydata.org; Waskom et al., 2016).

**Figure 2.**
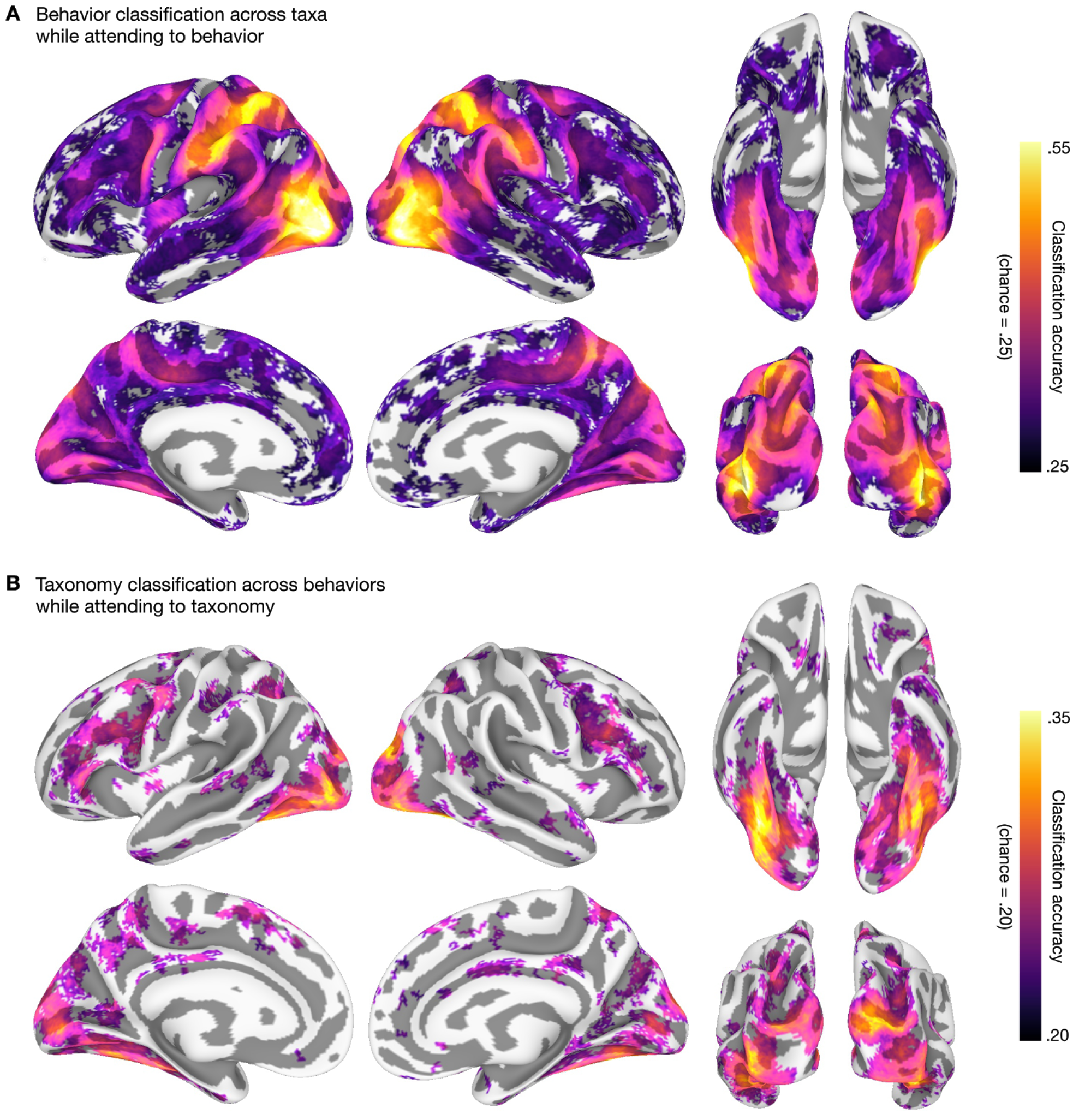
Behavioral and taxonomic category cross-classification using surface-based searchlights. To statistically evaluate the searchlight results, we first computed a one-sample t-test against theoretical chance accuracy per searchlight (one-tailed test). We corrected for multiple tests by controlling the false discovery rate (FDR) at *q* = .05 (Benjamini and Hochberg, 1995; Genovese et al., 2002). The mean classification accuracy across participants is plotted and searchlight maps are thresholded at FDR *q* = .05. **(A)** Searchlight classification of behavioral categories cross-validated across taxonomic categories while participants attended to animal behavior. Theoretical chance accuracy for four-way behavioral category classification is .25. The maximum mean searchlight accuracy for behavioral category classification was .56 in left lateral occipitotemporal cortex (inferior occipital gyrus). **(B)** Searchlight classification of taxonomic categories cross-validated across behavioral categories while participants attended to animal taxonomy. Theoretical chance accuracy for five-way taxonomic category classification is .20. The maximum mean searchlight accuracy for taxonomic category classification was .36 in right ventral temporal cortex (lateral fusiform gyrus). Although we used a *t*-test here for simplicity, note that the *t*-test may yield significant *t*-values even for near-chance accuracies, and a permutation- or prevalence-based approach may be preferable in some cases (cf. Allefeld et al., 2016; Etzel, 2017; Stelzer et al., 2013). Surface vertices on the medial wall were excluded from the analysis and clusters of fewer than ten contiguous significant vertices after thresholding were excluded for visualization purposes. Surface data were visualized using SUMA (Saad et al., 2004) and figures were created using GIMP (https://www.gimp.org) and Inkscape (https://inkscape.org).

## Conflict of Interest Statement

The authors declare that the research was conducted in the absence of any commercial or financial relationships that could be construed as a potential conflict of interest.

## Author Contributions

SAN, JVH, ACC, and MIG designed the experiment; SAN collected and analyzed the data; SAN, YOH, and JVH wrote the manuscript; SAN and YOH curated the data for public sharing.

## Funding

This work was supported by the National Institute of Mental Health at the National Institutes of Health (grant numbers F32MH085433-01A1 to ACC; and 5R01 MH075706 to JVH), and the National Science Foundation (grant numbers NSF1129764 and NSF1607845 to JVH).

## Acknowledgments

We thank Jason Gors, Kelsey G. Wheeler, Matteo Visconti di Oleggio Castello, J. Swaroop Guntupalli, Courtney Rogers, and Terry Sackett for assistance in collecting stimuli and data.

